# A Ferroptosis-mediated regulation of the biogenesis of the oxidative phosphorylation system

**DOI:** 10.1101/2022.02.17.480976

**Authors:** Anjaneyulu Murari, Shauna Kay-Rhooms, Kaniz BF Hossain, Tong Liu, Hong Li, Naga Sri Goparaju, Cindy Osei, Edward Owusu-Ansah

## Abstract

Several subunits in the matrix domain of mitochondrial complex I (CI) have been posited to be redox sensors for CI; but how elevated levels of reactive oxygen species (ROS) impinge on CI assembly is unknown. We report that when the mitochondrial NADPH-generating enzyme – Isocitrate Dehydrogenase 2 – is genetically disrupted, ROS levels are elevated and assembly of the oxidative phosphorylation system (OXPHOS) is impaired. Mechanistically, this begins with a ROS-mediated inhibition of biosynthesis of the matrix domain of CI, which progresses to a point where ferroptotic signals are induced, the mitochondrial unfolded protein response is activated and multiple OXPHOS complexes are impaired. Disruption of other enzymes that eliminate hydrogen peroxide, but not those that eliminate the superoxide radical, recapitulates the phenotype; implicating hydrogen peroxide as the signaling molecule involved. Thus, the redox status of the mitochondrion modulates the assembly of the matrix domain of CI and ultimately that of the entire OXPHOS.

## INTRODUCTION

Mammalian CI (NADH: ubiquinone oxidoreductase) contains 44 distinct subunits, one of which appears twice in the complex, giving rise to a total of 45 subunits. 14 subunits conserved from the ancestral enzyme in prokaryotes to eukaryotes constitute the minimal form of the enzyme. Consequently, they are referred to as the core or central subunits as they contain all the catalytic centers of the enzyme. The additional 30 unique subunits are referred to as accessory subunits. The 45 subunits are organized into two arms of the complex – referred to as the matrix and membrane domains – that are oriented almost orthogonally to each other, resulting in a boot- shaped structure [1]. There are 3 distinct functional modules of CI referred to as the N, Q and P- modules. The N-module is the site of NADH oxidation in the matrix domain and contains the FMN prosthetic group. The Q-module connects the N-module to the membrane domain; and functions as the conduit for electron transfer to Ubiquinone. The proton-pumping P-module is localized to the membrane domain (reviewed in [2–4]).

Several high-resolution cryoelectron micrographs have defined the locations of the accessory subunits in atomic detail; and shown that some have co-factors or features that make them excellent candidates for sensing the redox status of the cell [5–7]. For instance, the N-module subunit NDUFS6 interacts with Zn^2+^; and because Zn-containing proteins are very sensitive to oxidative stress, NDUFS6 has been proposed as a redox sensor. Similarly, a thioredoxin-like subunit, NDUFA2, also located in the N-module, has a pair of cysteine residues that are capable of forming disulfide bonds; giving rise to the hypothesis that NDUFA2 could also serve as a redox sensor for the complex. In addition, the Q-module subunit, NDUFA9, interacts with a molecule of NADPH making it a plausible sensor of NADPH-mediated redox alterations. Indeed, as Fe-S clusters as a group are very sensitive to inhibition by oxidative stress, any or all of the 8 Fe-S clusters in the matrix domain could in principle serve as redox sensors. Accordingly, we explored the link between redox homeostasis and CI assembly.

As a major source of ROS, mitochondria are equipped with a sophisticated antioxidant system that effectively buffers the organelle from the deleterious effects of ROS. In fact, mitochondria have co- opted moderate levels of ROS as signaling molecules that can be used to communicate to the nucleus to alter gene expression, in a phenomenon referred to as the mitochondrial integrated stress response [8, 9]. The FMN molecule in the N-module is the primary site where ROS is generated in CI. The initial ROS generated is the superoxide radical, which can be converted to hydrogen peroxide by the mitochondria-localized superoxide dismutase 2 (SOD2). There are also cytosolic and extracellular isoforms of SOD (i.e. SOD1 and SOD3 respectively).

Elimination of Hydrogen Peroxide represents a major step in ROS detoxification as it prevents the formation of the highly reactive hydroxyl radical, which can be formed when hydrogen peroxide reacts with Fe^2+^ in the Fenton reaction. Accordingly, hydrogen peroxide generated from the activity of the various SODs as well as other enzymes such as xanthine oxidase can be subsequently degraded into water by catalase or Glutathione Peroxidase. Glutathione Peroxidase uses reduced glutathione to eliminate hydrogen peroxide and lipid peroxides, in a process that converts reduced glutathione (GSH) to the oxidized form (GSSG). GSSG is recycled to GSH by glutathione reductase (GR) in a reaction that also converts NADPH to NADP^+^. Thus, NADPH-generating enzymes in the mitochondrion are a critical component of the mitochondrial antioxidant system. Finally, there are several peroxiredoxins and thioredoxins that also contribute to the antioxidant buffering capacity of the mitochondrion.

We previously showed that the mechanism of CI assembly in *Drosophila* flight (thoracic) muscles is similar to what has been described in mammalian systems [2, 10, 11]. This, together with its classical genetics and copious amounts of mitochondria, make the *Drosophila* flight muscles an ideal system for dissecting the mechanistic link between ROS formation and OXPHOS assembly *in vivo*. Accordingly, we knocked down or overexpressed several genes with antioxidant activity; and examined which aspect of OXPHOS assembly was impaired. We find that RNAi-mediated disruption of enzymes that eliminate hydrogen peroxide, but not those that eliminate the superoxide radical impair OXPHOS assembly. In particular, when the mitochondrial NADPH-generating enzyme, Isocitrate Dehydrogenase 2 (IDH2) is genetically disrupted, ROS levels are elevated and CI assembly is impaired. Mechanistically, this begins with a ROS-mediated inhibition of biosynthesis of the Q- and N-modules in the matrix domain of CI; that ultimately progresses to a point where pro-ferroptotic signals are activated, the Apoptosis Inducing Factor (AIF) is downregulated, and the mitochondrial unfolded protein response is upregulated. Notably, disruption of the mitochondrial NADP-dependent isoform of Malate Dehydrogenase recapitulates the phenotype. We conclude that a redox signaling axis involving mitochondrial NADPH-generating enzymes and ferroptosis activation modulates CI assembly, and ultimately the rest of the OXPHOS. This links CI assembly to the redox status of the mitochondrion.

## RESULTS

### RNAi-mediated disruption of the *Drosophila* ortholog of IDH2 (CG7176) and some antioxidant enzymes in *Drosophila* flight muscles impair OXPHOS assembly

The flight muscles in *Drosophila* are highly enriched with mitochondria and have been used extensively in the past to study the functions of proteins that regulate mitochondrial function [11–16]. Accordingly, to explore the possible role of ROS metabolism in OXPHOS assembly, we decided to examine the effect of knocking down or overexpressing various antioxidant enzymes on the assembly of OXPHOS complexes in flight muscles using the Gal4/UAS system [17]. Specifically, female flies carrying the Dmef2-Gal4 transgene were mated with various male flies each of which expressed either a UAS-RNAi transgene or UAS-cDNA transgene of a particular antioxidant enzyme; to obtain offspring that have a specific antioxidant enzyme either knocked down, or overexpressed respectively, in flight muscles (**Figure 1A**) [18].

**Figure 1:**
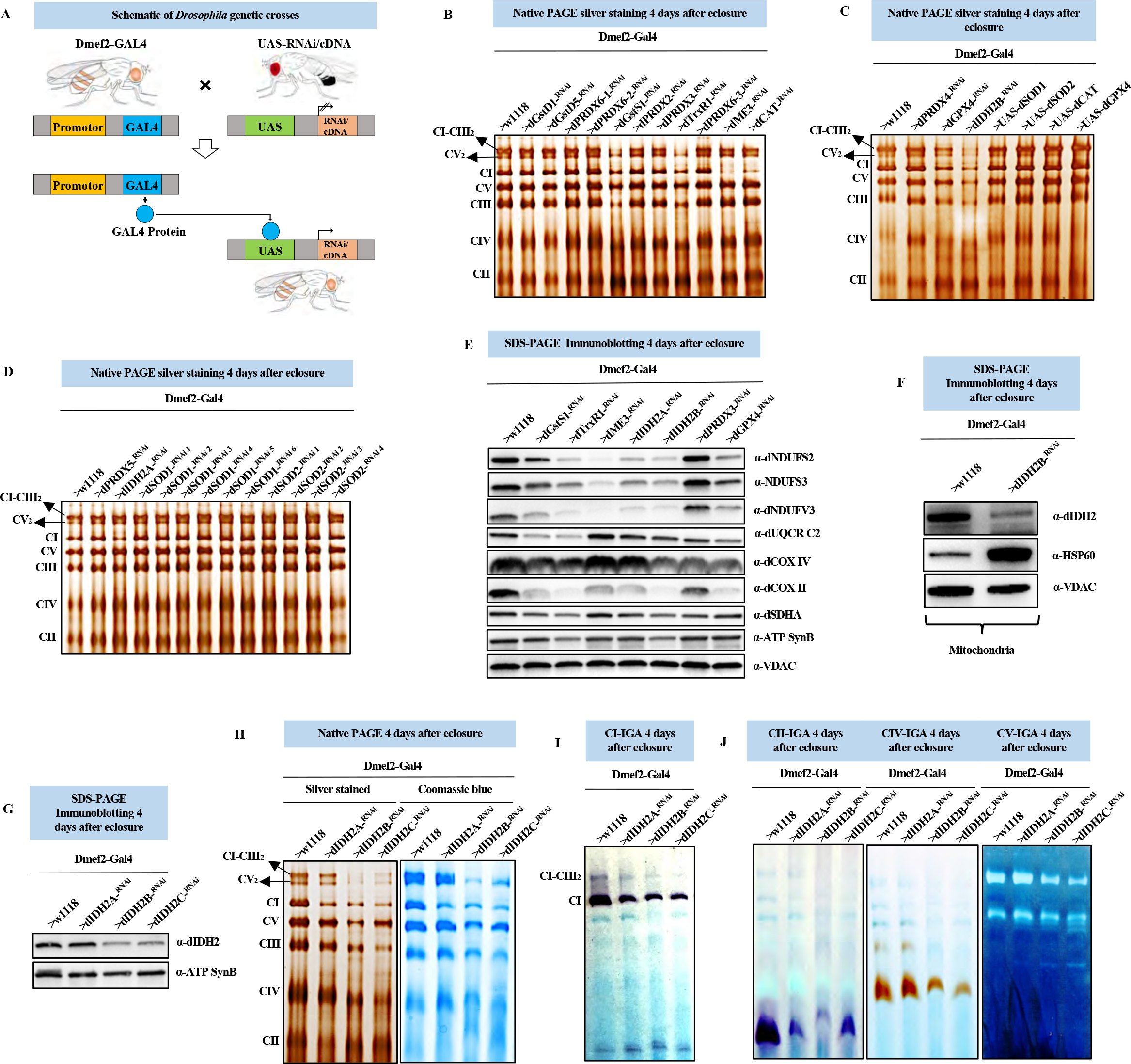
RNAi-mediated disruption of the *Drosophila* ortholog of IDH2 (CG7176) and some antioxidant enzymes in *Drosophila* flight muscles impair OXPHOS assembly. (A) A schematic describing the Gal4-UAS system. Temporal or tissue-specific expression of an RNAi or cDNA of interest is achieved in offspring produced from a cross between two transgenic flies: one carrying the RNAi or cDNA linked to an upstream activation sequence (i.e. UAS-RNAi or UAS-cDNA), and the other with the temporal or tissue-specific Gal4. In this manuscript Dmef-Gal4 (i.e. Gal4 is expressed under the control of the Dmef2 promoter in muscles) is used to express all RNAi or cDNA transgenes. Hence when the Dmef2-Gal4 flies are mated with the UAS-RNAi/cDNA flies, they will produce offspring where the RNAi or cDNA construct is expressed specifically in muscles. (B-D) Mitochondrial preparations from adult thoracic muscles expressing the UAS-RNAi or UAS- cDNA constructs shown in muscles (using Dmef2-Gal4) 4 days after eclosure, were analyzed by silver staining of native gels (see methods). (E) Total cell lysates from flight/thoracic muscles isolated from Dmef2-Gal4/w1118 (wildtype) and other flies expressing the UAS-RNAi constructs shown with the Dmef2-Gal4 transgene 4 days after eclosure, were analyzed by SDS-PAGE and immunoblotting with anti-dNDUFS2, anti- NDUFS3, anti-dNDUFV3, anti-dUQCRC2, anti-dCOXIV, anti-dCOXII, anti-dSDHA and Anti-ATP Synthase Beta (ATP5F1B). Anti-VDAC was used as a loading control. (F) Mitochondrial preparations from Dmef2-Gal4/w1118 (wildtype) and Dmef2-Gal4/UAS-dIDH2B^- RNAi^ adult thoracic muscles, were analyzed by SDS-PAGE and immunoblotting with anti-dIDH2 and anti-Hsp60. Anti-VDAC was used as a loading control. (G) Total cell lysates from flight/thoracic muscles isolated from Dmef2-Gal4/w1118 (wildtype) and other flies expressing the 3 different UAS-RNAi constructs to dIDH2 with the Dmef2-Gal4 driver 4 days after eclosure, were analyzed by SDS-PAGE and immunoblotting with anti-dIDH2. Anti-ATP Synthase Beta (ATP5F1B) was used as a loading control. (H) Mitochondrial preparations from adult thoracic muscles expressing the transgenic RNAi constructs shown 2 days after eclosure, were analyzed by silver staining of native gels and blue native polyacrylamide gel electrophoresis (BN-PAGE). The asterisk (*) denotes accumulated assembly intermediates. (I and J) Complex I, complex II, complex IV and complex V in-gel activity assays of OXPHOS complexes obtained from mitochondrial preparations from flight muscles of flies with the genotypes shown.

We knocked down the expression of genes encoding for *Drosophila* orthologs of Superoxide Dismutase 1 (*sod1*), Superoxide Dismutase 2 (*sod2*), Catalase (*cat*), Isocitrate Dehydrogenases 2 (*idh2*), the NADPH-dependent Malate Dehydrogenase (*men-b;* hereafter referred to as dME3), multiple Peroxiredoxins (*prdx2, prdx3, prdx4, prdx5, prx6005/prdx6-1, prx2540-1/prdx6-2,* and *prx2540-2/prdx6-3*), multiple Glutathione S-Transferases (*gstD1, gstD5* and *gstS1*), Thioredoxin 1 (*trxr-1*), and Glutathione Peroxidase 4 (*gpx4*) in flight muscles (see methods). Subsequently, we isolated mitochondria from the thoraxes of adult flies expressing the various RNAi constructs, solubilized their membranes in digitonin, and analyzed the integrity of their OXPHOS complexes using silver staining of blue native gels (**Figures 1B-1D**). RNAi-mediated disruption of GstS1, Trxr- 1, GPX4 and one of the transgenic RNAi constructs targeting IDH2 impaired the assembly of multiple OXPHOS complexes, but especially CI and CIII (**Figures 1B-1D**). On the other hand, RNAi-mediated knockdown of dME3, Cat, and a second RNAi construct for IDH2 impaired CI assembly (**Figures 1B-1D**). Surprisingly, none of the multiple transgenic RNAi lines targeting *sod1* and *sod2*, including a few that were potent enough to cause the flies to succumb to lethality within 3 days of eclosing as adults, notably impaired OXPHOS assembly (**Figure 1D**). In addition, none of the antioxidant enzymes we overexpressed (i.e. Sod1, Sod2, Cat and GPX4) disrupted OXPHOS assembly (**Figure 1C)**. Immunoblotting of OXPHOS complexes from samples where the RNAi lines showed phenotypes, confirmed the silver staining results **(Figure 1E)**. These results indicate that a failure to eliminate the superoxide radical while deleterious, does not appear to alter OXPHOS assembly. Indeed, all the enzymes that produced an assembly defect when disrupted are required for eliminating hydrogen peroxide or lipid peroxides.

Isocitrate Dehydrogenases (IDH) catalyze the oxidative decarboxylation of isocitrate to 2- oxyglutarate and a reduced pyridine nucleotide cofactor. Three major isoforms of IDH can be distinguished based on their subcellular localization and type of pyridine nucleotide co-factor used to catalyze their enzymatic reaction. IDH1, localized to the cytosol and peroxisomes, and IDH2 which is localized to the mitochondrial matrix, use nicotinamide adenine dinucleotide phosphate (NADP^+^) as cofactor. IDH3 which is also localized to the mitochondrial matrix but uses nicotinamide adenine dinucleotide (NAD^+^) as co-factor, is a component of the TCA cycle. CG7176 is the sole *Drosophila* ortholog of both IDH1 and IDH2, suggesting that it could function in the cytosol, peroxisome and/or mitochondrion. We used an antibody we raised against CG7176 to confirm that at least a portion of CG7176 localizes to the mitochondrion (**Figure 1F**). We refer to CG7176 as dIDH2 in this manuscript, and prefix all *Drosophila* orthologs of CI subunits with “d” (as in dNDUFS3 for NDUFS3, etc) for the rest of this manuscript.

We knocked down IDH2 expression in flight muscles using three different transgenic RNAi constructs, and compared the integrity of their OXPHOS complexes using silver staining and blue native PAGE (BN-PAGE). The genotypes of the flies were *Dmef2-Gal4/UAS-dIDH2^-RNAi-1^* (dIDH2- A), *Dmef2-Gal4/UAS-dIDH2^-RNAi-2^* (dIDH2-B), and *Dmef2-Gal4/UAS-dIDH2^-RNAi-3^* (dIDH2-C).

Immunoblotting analyses revealed that the expression of dIDH2 had been notably reduced in thoraces obtained from dIDH2-B and dIDH2-C flies (**Figure 1G)**. In contrast to the specific CI assembly defect observed in dIDH2-A thoraces, the assembly of multiple complexes were impaired in mitochondria from dIDH2-B and dIDH2-C flies (**Figure 1H).** In line with the silver staining and BN-PAGE results described in Figure 1H, in gel CI activity was reduced in all three genotypes (**Figure 1I**). In gel CII, CIV and CV activity were also reduced in mitochondria from dIDH2-B and dIDH2-C flies (**Figure 1J**). Taken together, these observations establish the order of severity of phenotypes produced by the three transgenic RNAi lines used as: dIDH2-A (mild), dIDH2-B (intermediate) and dIDH2-C (severe). They also show that dIDH2 is a major regulator of OXPHOS assembly. Therefore, the three transgenic RNAi constructs provide an opportunity to explore the progressive nature of OXPHOS damage triggered by the loss of IDH2 in the adult thoracic muscles of *Drosophila*.

### RNAi-mediated disruption of dIDH2 activates ferroptotic signals

To explore the functional consequences of disrupting dIDH2 expression, we examined the effect of knocking down dIDH2 in flight muscles, on the locomotory activity of the flies. Overt climbing defects were observed in the dIDH2-B flies (**Figure 2A**). To further validate the locomotory defects observed in Figure 2A, we monitored the spontaneous locomotory activity of dIDH2-A and dIDH2- B flies relative to wildtype flies (*Dmef2-Gal4/w1118)* within the first two weeks after the flies eclosed as adults. When analyzed at 25C, the spontaneous locomotory ability of dIDH2-B flies were markedly impaired starting around 5 days (120 hours) after eclosure; but this was less pronounced for the dIDH2-A flies that failed to produce a readily perceptible climbing defect (**Figures 2A and 2B)**. The NADP: NADPH ratio was increased in thoraces from dIDH2-A and dIDH2-B flies, in agreement with the established function of IDH2 in reducing NADP^+^ to NADPH (**Figure 2C**). An Amplex Red assay revealed that there was an increase in the amount of Hydrogen Peroxide - an indication of oxidative stress - in both dIDH2-A and dIDH2-B samples, relative to wildtype controls (**Figure 2D**). Increased oxidative stress can cause lipid peroxidation, culminating in the formation of thiobarbituric acid reactive substances (TBARS), which can be detected by a TBARS assay (see methods). The TBARS assay revealed that the extent of lipid peroxidation was increased in dIDH2-B samples (**Figure 2E**). However, in spite of these findings, caspase activity was decreased in thoraces from dIDH2-A and dIDH2-B flies (**Figure 2F**).

**Figure 2:**
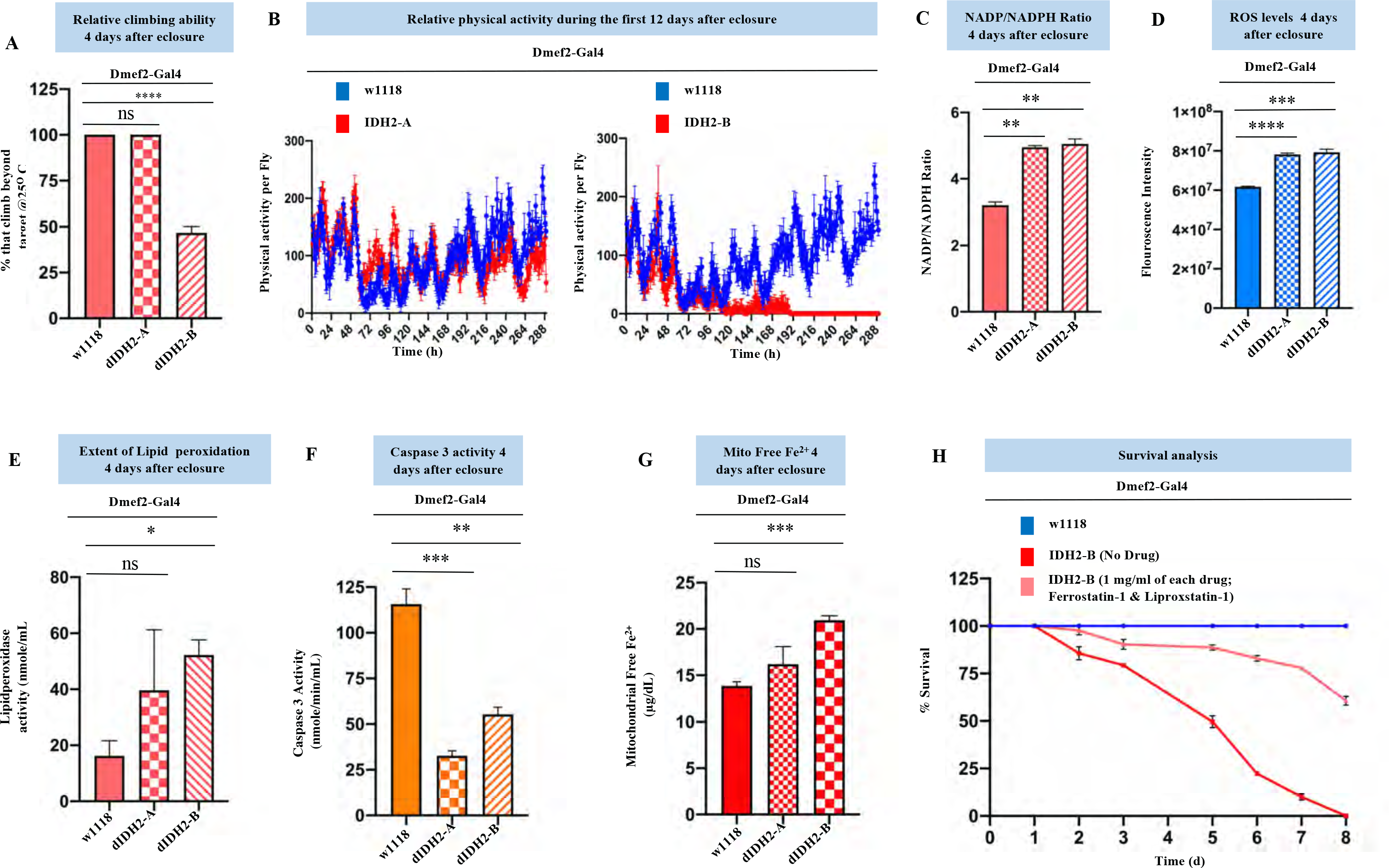
RNAi-mediated disruption of dIDH2 activates ferroptotic signals. (A) Relative climbing ability of *Dmef2-Gal4/UAS-dIDH2^-RNAi-1^* (dIDH2-A), *Dmef2-Gal4/UAS-dIDH2^- RNAi-2^* (dIDH2-B) and *Dmef2-Gal4/w1118 (w1118,* wildtype controls) 4 days after eclosure. (B) Relative spontaneous physical activities of dIDH2-A, dIDH2-B and wildtype flies during the first 12 days after eclosure (C) NADP:NADPH ratio of dIDH2-A, dIDH2-B and wildtype flies 4 days after eclosure. (D) Relative ROS levels of dIDH2-A, dIDH2-B and wildtype flies 4 days after eclosure as determined by an Amplex Red Assay (E) Lipid peroxidation assay of dIDH2-A, dIDH2-B and wildtype flies 4 days after eclosure (F) Relative caspase 3 activity levels of dIDH2-A, dIDH2-B and wildtype flies 4 days post-eclosure (G) Relative mitochondrial free Ferrous (Fe^2+^) levels of dIDH2-A, dIDH2-B and wildtype flies 4 days post-eclosure. (H) Survival curves of dIDH2-B, wildtype flies, and dIDH2-B flies raised on a diet supplemented with 1mg/ml each of two Ferroptosis inhibitors (Ferrostatin-1 and Liproxstatin-1). In all instances, n = 3 biological replicates with 100 flies per replicate; p values are based on the student’s t-test for unpaired two-tailed samples. The fold change shown refers to the mean ± s.e.m (standard error of the mean); and n.s. denotes p > 0.05, * = p<0.05, ** = p<0.01 and *** = p<0.001.

Ferroptotic cell death is a non-apoptotic form of cell death that is triggered by iron-dependent lipid peroxidation [19]. As the extent of lipid peroxidation was elevated, but caspase activity was reduced in thoraces from dIDH2-B flies, we wondered whether ferroptosis was induced under these conditions. To this end, we assessed and found that the amount of labile ferrous iron (free Fe^2+^) was increased in the mitochondrion of dIDH2-B flies relative to wildtype controls (**Figure 2G**). Remarkably, raising the dIDH2-B flies on a diet supplemented with two inhibitors of lipid peroxidation – Ferrostatin-1 and Liproxstatin-1 – potently rescued the early lethality of dIDH2-B flies (**Figure 2H**). Thus, an integration of the bioenergetic and functional data shown in Figures 1 and 2 indicate that disruption of dIDH2 causes an upregulation of several pro-ferroptotic signals to impair OXPHOS assembly and elicit ferroptotic cell death. Consequently, we pursued additional studies to uncover the molecular mechanism by which the ferroptotic-mediated disruption of dIDH2 regulates OXPHOS assembly.

### The synthesis of matrix-localized CI assembly intermediates are impaired when dIDH2 is disrupted

Mitochondrial CI has a characteristic L-shaped structure. It consists of a matrix domain extending into the mitochondrial matrix, oriented almost perpendicularly to a membrane domain localized to the mitochondrial inner membrane (**Figure 3A**) [20, 21].

**Figure 3:**
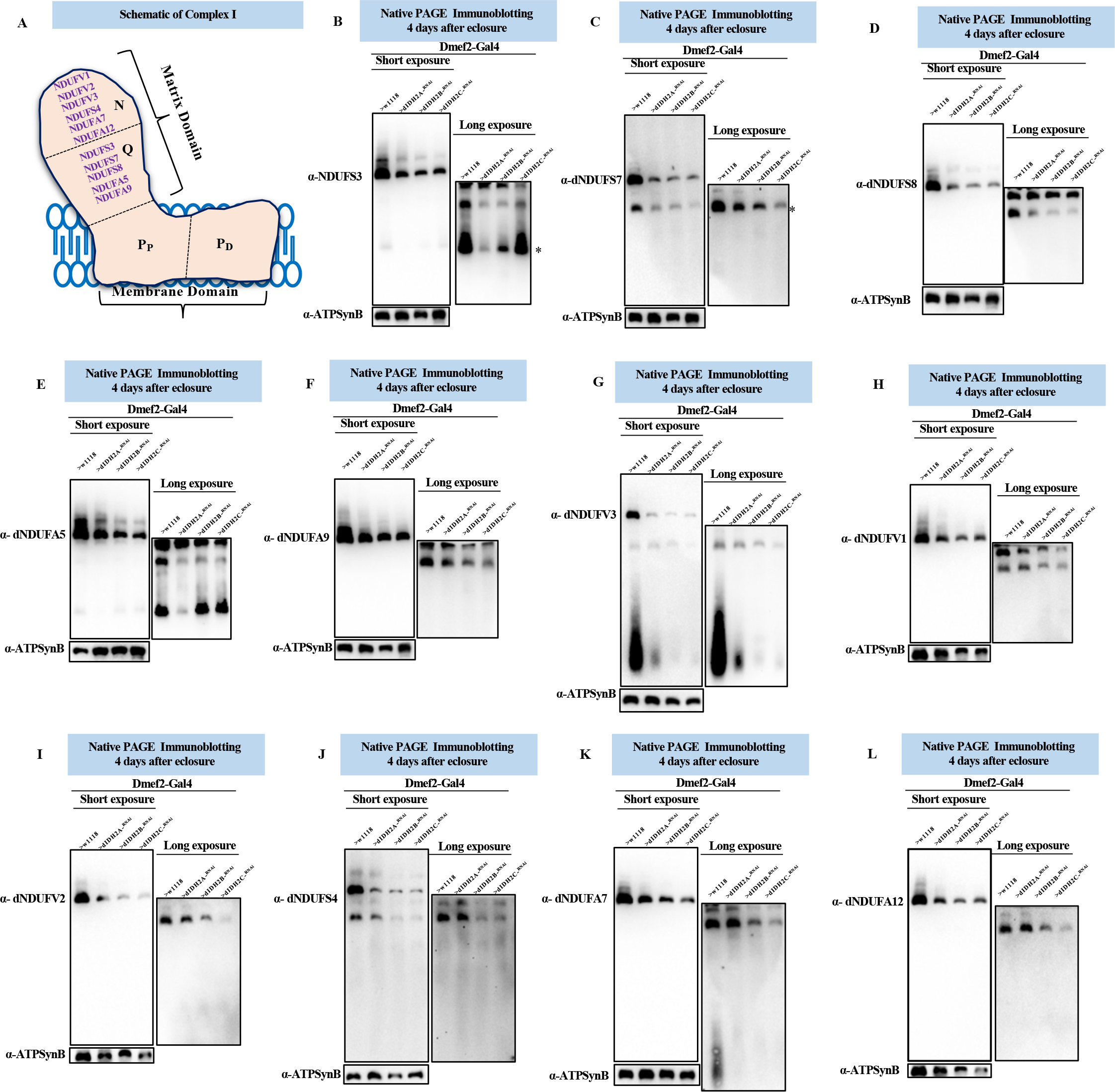
The synthesis of matrix-localized CI assembly intermediates are impaired when dIDH2 is disrupted. (A) An illustration of mitochondrial CI depicting the approximate positions of the N, Q, PP and PD modules. The matrix and membrane domains are oriented almost orthogonally to each other, resulting in a boot-shaped structure. A commercially available antibody that detects dNDUFS3, and antibodies we raised against dNDUFA5, dNDUFA9, dNDUFS7 and dNDUFS8 were used to examine the biogenesis of the Q-module. Biogenesis of the N-module was tracked by immunoblotting with antibodies we raised against dNDUFS4, dNDUFA7, dNDUFA12, dNDUFV1, dNDUFV2 and dNDUFV3. (B-L) Mitochondrial preparations from thoraces isolated from Dmef2-Gal4/w1118 (wildtype), dIDH2-A, dIDH2-B and dIDH2-C flies 4 days post-eclosure were analyzed by BN-PAGE, followed by immunoblotting with the antibodies indicated. The blots were imaged following a short exposure to detect the holoenzyme and supercomplexes, after which the region corresponding to the holoenzyme and supercomplexes was cut off, and the rest of the blot re-imaged after a longer exposure to detect the assembly intermediates. The antibodies used were anti-NDUFS3 which detects dNDUFS3 (B), anti-dNDUFS7 (C), anti-dNDUFS8 (D), dNDUFA5 (E), anti-dNDUFA9 (F), anti-dNDUFV3 (G), anti-dNDUFV1 (H), dNDUFV2 (I), anti-dNDUFS4 (J), anti-dNDUFA7 (K), and anti-dNDUFA12 (L). Anti-ATP Synthase Beta (ATP5F1B) which detects the *Drosophila* ortholog, dATP-Syn*β* was used as a loading control.

To begin to uncover the mechanism by which dIDH2 regulates OXPHOS assembly, we examined the effect of disrupting dIDH2 on the biogenesis of the matrix domain of CI (**Figure 3A**). During CI assembly, specific assembly intermediates (AIs) consisting of a few CI subunits form largely independently of each other and merge in a stereotypic fashion en route to forming the mature complex [3, 10, 22]. The N-, Q- and P-modules are synthesized from specific AIs.

Synthesis of the Q-module begins with the formation of an AI consisting of dNDUFS2 and dNDUFS3, which ultimately combines with dNDUFS7, dNDUFS8, dNDUFA5 and dNDUFA9. Consequently, we tracked the biogenesis of the Q-module *via* immunoblotting of blue native gels with antibodies that detect dNDUFS3, dNDUFS7, dNDUFS8, dNDUFA5 and dNDUFA9. There was a reduction in the amount of dNDUFS3 in an initiating AI of the Q-module in mitochondria isolated from thoraxes of dIDH2-A, dIDH2-B and dIDH2-C flies. Similarly, the amount of dNDUFS7, dNDUFS8, dNDUFA5 and dNDUFA9 that had incorporated into the Q-module was diminished in all three RNAi backgrounds. (**Figures 3B-3F**).

The N-module consists of the following subunits: dNDUFV1, dNDUFV2, dNDUFV3, dNDUFA2, dNDUFA6, dNDUFA7, dNDUFA12, dNDUFS1, dNDUFS4 and dNDUFS6. Accordingly, we tracked the incorporation of dNDUFV1, dNDUFV2, dNDUFV3, dNDUFS4, dNDUFA7 and dNDUFA12 into CI AIs to monitor the integrity of the N module. We observed that the stabilization or incorporation of dNDUFV3 into an initiating AI of the N-module was impaired in mitochondria isolated from thoraxes of dIDH2-A, dIDH2-B and dIDH2-C flies (**Figure 3G**). This was coupled with a decrease in the amount of dNDUFV1, dNDUFV2, dNDUFS4, dNDUFA7 and dNDUFA12 that had incorporated into subcomplexes of the N-module isolated from mitochondria obtained from all three RNAi backgrounds; but most prominently from the dIDH2-B and dIDH2-C flies (**Figures 3H-3L**). Taken together, we conclude that disruption of IDH2 impairs the biogenesis or stability of the Q and N-modules.

### dIDH2 regulates the biogenesis of some assembly intermediates in the mitochondrial inner membrane

The membrane domain is comprised of the proton-pumping P-module; which can be further subdivided into a proximal PP module and a distal PD module. 5 of the 7 mt-DNA-encoded subunits (i.e. dND1, dND2, dND3, dND4L, and dND6) are part of the PP module, while dND4 and dND5 are localized to the PD module (**Figure 4A**) (reviewed in [3, 23]).

**Figure 4:**
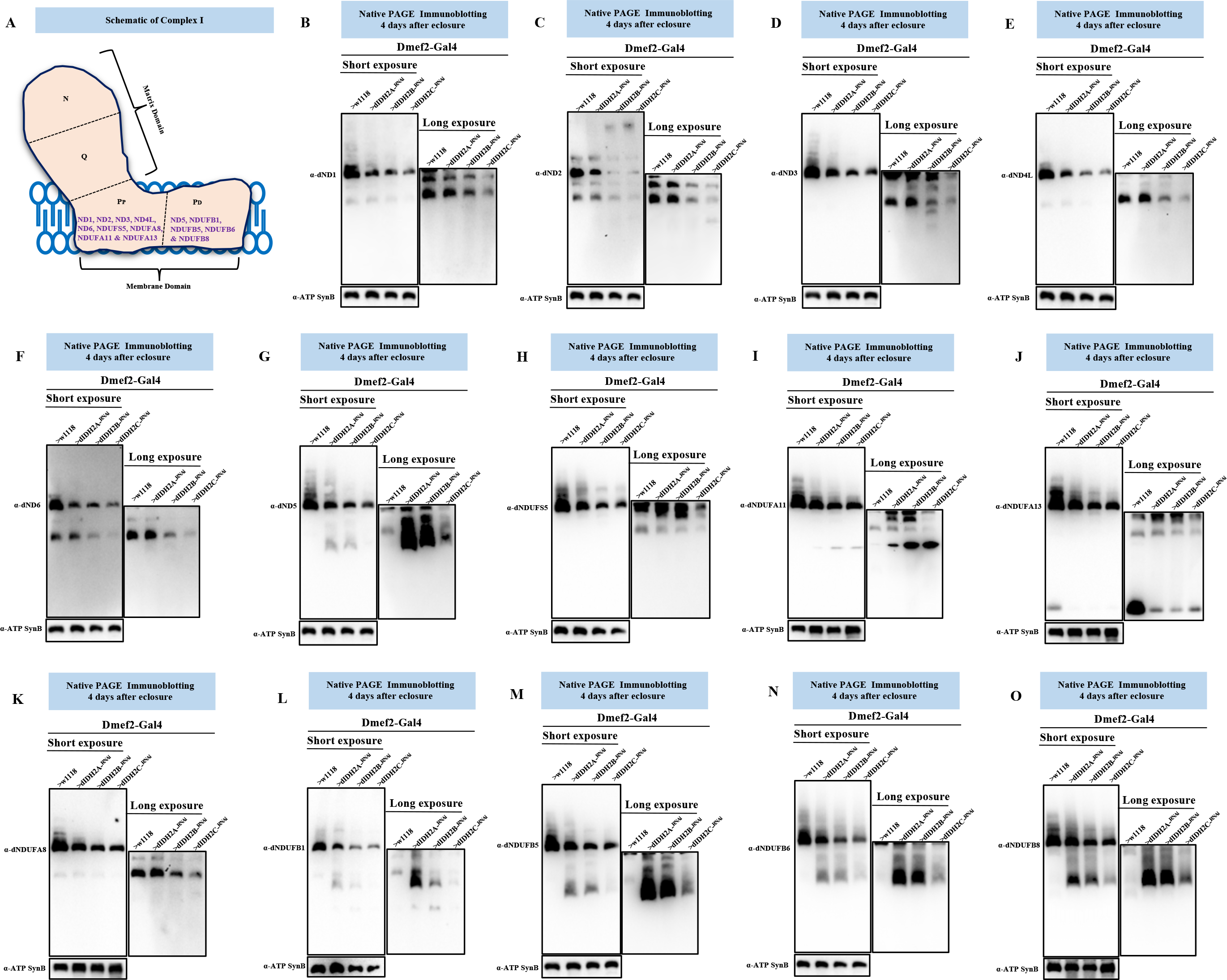
dIDH2 regulates the biogenesis of some assembly intermediates in the mitochondrial inner membrane. (A) A diagram of mitochondrial CI showing the approximate positions of the N, Q and P modules. The P module can be sub-divided into the PP and PD sub-modules. Antibodies raised against the subunits shown were used to monitor the synthesis of the PP and PD sub-modules. (B-O) Mitochondrial preparations from thoraces isolated from Dmef2-Gal4/w1118 (wildtype), dIDH2-A, dIDH2-B and dIDH2-C flies 4 days after eclosure were analyzed by BN-PAGE, followed by immunoblotting with the antibodies listed. The blots were imaged following a short exposure to detect the holoenzyme and supercomplexes, after which the region corresponding to the holoenzyme and supercomplexes was cut off, and the rest of the blot re-imaged after a longer exposure to detect the assembly intermediates. The antibodies used were anti-dND1 (B), anti- dND2 (C), anti-dND3 (D), anti-dND4L (E), anti-dND6 (F), anti-dND5 (G), dNDUFS5 (H), anti- dNDUFA11 (I), anti-dNDUFA13 (J), anti-dNDUFA8 (K), anti-dNDUFB1 (L), anti-dNDUFB5 (M), anti-dNDUFB6 (N),and anti-dNDUFB8 (O). Anti-ATP Synthase Beta (ATP5F1B) which detects the *Drosophila* ortholog, dATP-Syn*β* was used as a loading control.

We explored whether disruption of dIDH2 impairs the incorporation of the mt-DNA-encoded CI subunits into the P-module (**Figure 4A**). The incorporation of dND1 into the PP module of samples from dIDH2-A was slightly disrupted, but was progressively more impaired in dIDH2-B and dIDH2- C samples respectively (**Figure 4B**). Similarly, incorporation of dND2 was not appreciably perturbed in dIDH2-A thoraces, but showed potent reductions in dIDH2-B and dIDH2-C samples (**Figure 4C**). We previously showed that dND3, dND4L and dND6 are very unstable and are rapidly degraded when CI biogenesis is impaired [11]. Interestingly, a dND3-containing AI was somewhat stalled in dIDH2-A samples; and we observed what appeared to be a smear of the dND3- containing AI in dIDH2-B samples, indicating that the AI had stalled and was undergoing degradation (**Figure 4D**). dND4L- and dND6-containing AIs were also stalled in dIDH2-A, but were progressively more reduced in dIDH2-B and dIDH2-C samples respectively (**Figures 4E and 4F**). Finally, a dND5-containing AI (part of the PD module) was robustly backlogged in dIDH2-A and dIDH2-B samples, but not in dIDH2-C samples; although there appeared to be remnants of a dND5-containing AI that had degraded in the dIDH2-C samples (**Figure 4G**).

We also examined the effect of knocking down dIDH2 expression on the incorporation of nuclear-encoded CI subunits into the P-module (**Figure 4A**). Incorporation of dNDUFS5 and dNDUFA11 appeared to accumulate in a terminal AI that migrates just below the CI holoenzyme in dIDH2-A and dIDH2-B thoraces; but the amount in dIDH2-C samples was reduced (**Figure 4H and 4I**). Similarly, immunoblotting with anti-dNDUFA13 detected the stalled terminal AI in dIDH2-A and dIDH2-B thoraces, but not in the dIDH2-C sample (**Figure 4J**). Furthermore, incorporation of dNDUFA8 was stalled in dIDH2-A, but the amount of the dNDUFA8-containing AI was decreased in dIDH2-B and dIDH2-C samples respectively (**Figures 4K**). In line with observations described in Figure 4G, immunoblotting against multiple PD module subunits (i.e. dNDUFB1, dNDUFB5, dNDUFB6 and dNDUFB8) showed that the PD module was severely backlogged in dIDH2-A and dIDH2-B samples (**Figure 4L-O**). In essentially all instances where a P-module AI accumulated in dIDH2-A and dIDH2-B thoraces but not in dIDH2-C samples, there appeared to be vestiges of the accumulated AI in dIDH2-C; which may be indicative of an initial accumulation of the AI in dIDH2-C that eventually succumbed to proteolytic degradation (**Figures 4G and 4L-O**). Collectively, the results described herein point to the following conclusion: there was a stalling and accumulation of several P-module AIs that was readily apparent in the dIDH2-A samples, but was sometimes observed in the dIDH2-B and dIDH2-C samples as well. In some instances, some subunits in the accumulated AI were rapidly degraded; an observation that was most noticeable in dIDH2-C samples. When combined with results from Figure 3, we infer that biogenesis of the P-module was not inhibited. However, the diminution in synthesis or stability of the Q-module when dIDH2 is genetically impaired causes a stalling and accumulation of multiple AIs in the P-module, which ultimately succumb to degradation.

### A robust compensatory adaptive response is induced as a result of genetic disruption of dIDH2

We employed a proteomics approach to gain further insight into the mechanism by which disruption of dIDH2 impacts the OXPHOS. Specifically, we isolated mitochondria from dIDH2-A and wildtype thoraxes, solubilized the mitochondrial membranes in digitonin and resolved their OXPHOS complexes by BN-PAGE. Subsequently, we excised a portion of the blue native gel that encompasses the CI holoenzyme, (hereafter referred to as profile A), extracted the constituent proteins, and analyzed them by mass spectrometry (**Figures 5A**). We used label-free spectral counting to assess the relative abundance of individual subunits in the 4 CI modules of the holoenzyme from dIDH2-A thoraxes relative to the wildtype sample [24–26].

**Figure 5:**
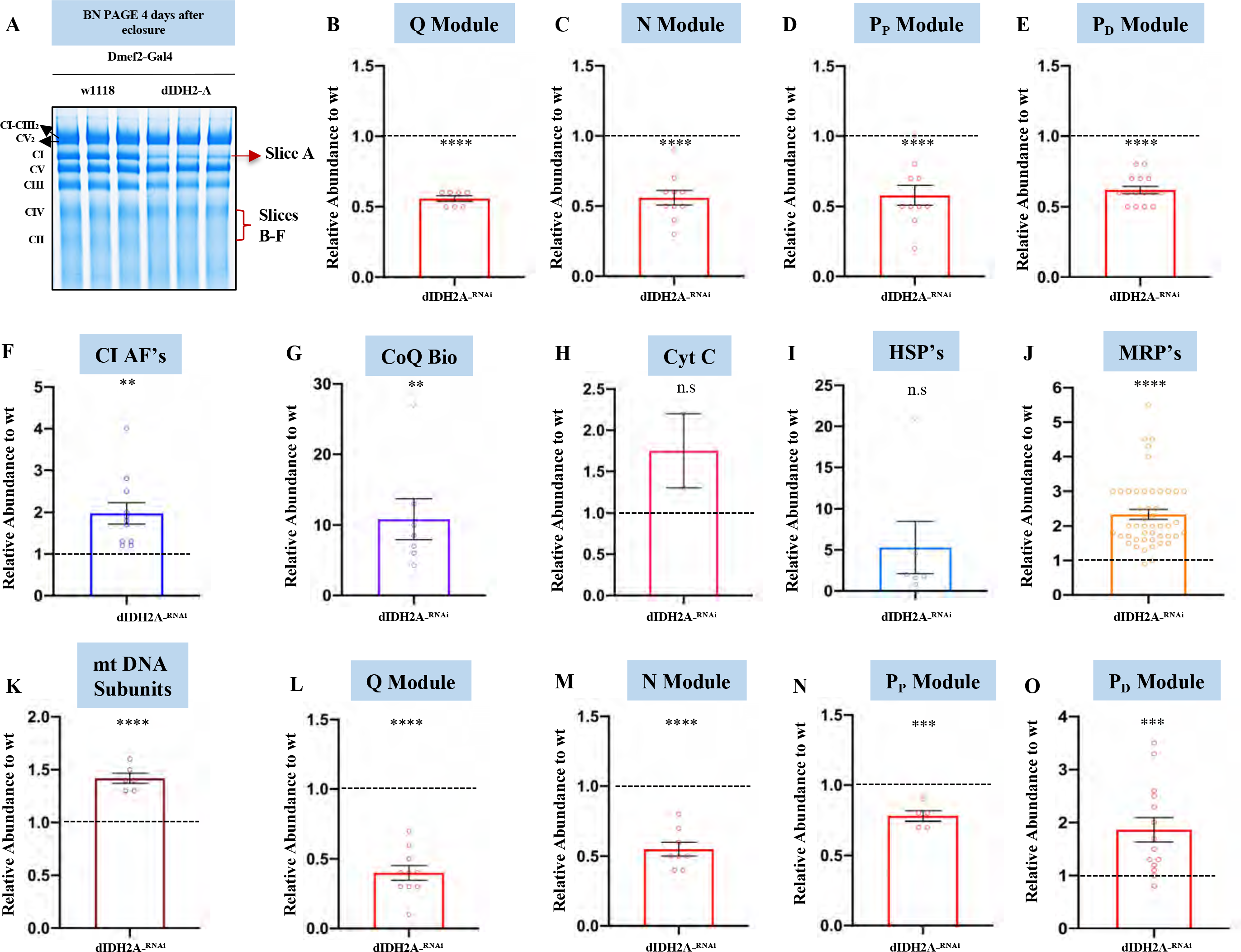
A robust compensatory adaptive response is induced as a result of genetic disruption of dIDH2. (A) A BN-PAGE gel showing the gel slices that were excised and analyzed by mass spectrometry. (B-E) Label-free quantitative proteomics was used to assess the relative amount of CI subunits in the holoenzyme (gel slice A) that are part of the Q module (B), N module (C), the PP module (D) and the PD module (E) in mitochondrial preparations from Dmef2-Gal4/w1118 (wildtype) and dIDH2-A fly thoraces. (F-K) Label-free quantitative proteomics was used to quantify the relative amount of CIAFs (F), genes that regulate Coenzyme Q biosynthesis (G), cytochrome C (H), mitochondrial heat shock proteins (I), mitochondrial ribosomal proteins (J), and mt-DNA-encoded OXPHOS subunits (K), that co-migrate with the holoenzyme (gel slice A) in mitochondrial preparations from Dmef2- Gal4/w1118 (wildtype) and dIDH2-A fly thoraces. (L-O) Label-free quantitative proteomics was used to assess the relative amount of CI subunits in the gel slice B (overlaps with CIV) that are part of the Q module (L), N module (M), the PP module (N) and the PD module (O) in mitochondrial preparations from Dmef2-Gal4/w1118 (wildtype) and dIDH2-A fly thoraces. In all 6 gel slices, n = 2 biological replicates with mitochondria from 20 fly thoraces per replicate; p values are based on the student’s t-test for unpaired two-tailed samples. The fold change shown refers to the mean ± s.e.m (standard error of the mean); and n.s. denotes p > 0.05, * = p<0.05, ** = p<0.01 and *** = p<0.001.

CI subunits in the Q, N, PP and PD modules were all downregulated in the gel slice corresponding to the holoenzyme; in line with observations from Figure 1 showing that CI assembly is impaired in thoraxes from dIDH2 flies (**Figures 5B-5E**, and **Table S1**). Several CI assembly factors (CIAFs) remained in association with the residual CI in the dIDH2-A samples (**Figure 5F and Table S1**). As many CIAFs function as chaperones to stabilize relatively unstable CI assembly intermediates, their association with the CI that remains in dIDH2-A thoraxes appears to be an adaptive response to preserve a sub-stoichiometric amount of CI. Notably, there was a robust upregulation of multiple proteins involved in co-enzyme Q biosynthesis that co-migrated with the residual CI in dIDH2-A thoraxes; and the two paralogs of cytochrome c in *Drosophila* were also upregulated in association with CI from dIDH2-A flight muscle mitochondria (**Figures 5G, 5H and Table S1**). These results are consistent with an adaptive response in mitochondria from dIDH2-A thoraxes aimed at ensuring the continuous passage of electrons through the electron transport chain in spite of a suboptimal CI. Some mitochondrial chaperones such as Hsp60 and Hsc70-5, both of which are components of the mitochondrial unfolded protein response were robustly upregulated in the vicinity of CI from dIDH2-A thoraxes (**Figures 5I and Table S1**). Additionally, there was a modest but consistent increase in the amount of several mitochondrial ribosomal proteins co-migrating with CI in dIDH2-A samples (**Figure 5J and Table S1**). As translation of mtDNA-encoded CI subunits and subsequent integration into the CI biosynthetic pathway occurs almost concurrently, these observations are consistent with the induction of a robust compensatory mitochondrial translation response aimed at forestalling CI biogenesis under sub-optimal conditions. Indeed, all other mtDNA-encoded OXPHOS subunits (mt: cyt-b in CIII; mt: coI, mt: coII, and mt: coIII in CIV; and mt: ATPase 6 and mt: ATPase 8 in CV) were upregulated (**Figure 5K and Table S1**). These results indicate that a robust compensatory adaptive response is induced in dIDH2-A thoraxes.

We previously showed that most of the initiating AIs migrate at a region between CII and CIV in blue native gels [10]. Therefore, we divided the region of the blue native gel between CII and CIV into 5 evenly-sized slices, labelled slice B to F, from higher to lower molecular weight and analyzed their proteomic constituents by mass spectrometry (**Figure 5A**). Multiple Q-, N-, and Pp-module subunits, as well as the CIAFs – NDUFAF3 and NDUFAF4 – were downregulated in the dIDH2-A sample in profile B (**Figure 5L-5N, and Table S2**). However, most PD module subunits were upregulated in profile B (**Figure 5O and Table S2**). Altogether, the wave of CI biogenesis observed in profile B indicates that initiating Q-, N-, and Pp-module AIs were unstable in the dIDH2-A sample resulting in a stalling and accumulation of the PD-module; as the PD-module can only progress further in the CI biosynthesis pathway if there are sufficient “binding partners” of the PP-module. Finally, we observed that in all profiles analyzed, genes that regulate mitochondrial protein homeostasis were upregulated in the dIDH2-A sample (**Tables S1-S6**). Altogether, these data reinforce our conclusion that a compensatory adaptive response aimed at repairing or degrading misfolded proteins in the mitochondrion is induced when dIDH2 is disrupted.

### Induction of the alternative NADH-ubiquinone oxidoreductase (*Ndi1)* partially restores the biogenesis of the Q-module and suppresses the extent of activation of the JNK and MAPK pathways

The alternative NADH-ubiquinone oxidoreductase (*Ndi1)* is a nuclear-encoded polypeptide in yeast that transfers electrons to Ubiquinone. In this respect, it compensates for the lack of a multi-subunit CI in yeast; although it lacks the proton-pumping ability of CI. Overexpression of the NDI1 enzyme in *Drosophila* leads to an increase in NADH–ubiquinone oxidoreductase activity [27]. Because of the degeneration of the matrix domain of CI when dIDH2 is knocked down, electrons donated to CI from NADH2, which under normal conditions are accepted by the FMN site in the N-module and subsequently transferred through the Q- module to Ubiquinone, are likely to stall in the vicinity of CI. Accordingly, we hypothesized that overexpression of NDI1 will suppress, at least some aspects of the phenotypes observed in dIDH2-B flight muscles.

To test this hypothesis, we obtained mitochondria from dIDH2-B flight muscles/thoraces overexpressing either GFP (UAS-GFP, as a negative control) or NDI1 (UAS-NDI1) with the Dmef2- Gal4 transgene, and evaluated their matrix domain CI AI profiles. In similarity to our findings reported in Figure 3B, biogenesis or stability of the Q-module, assessed by immunoblotting of dNDUFS3, was impaired in dIDH2-B/UAS-GFP flight muscles. However, overexpression of NDI1 in dIDH2-B flight muscles led to a notable restoration of the dNDUFS3-containing initiating AI (**Figure 6A**). NDUFA9 is an accessory subunit in the matrix domain that contains a tightly-bound NADPH molecule. In spite of the fact that biogenesis of the initiating dNDUFS3-containing AI is rescued as a result of NDI1 expression, the diminished incorporation of dNDUFA9 into the matrix domain cannot be ameliorated by overexpressing NDI1 (**Figure 6B**). This is likely due to the fact that although expression of NDI1 reduced the amount of ROS formed, it was incapable of restoring NADPH levels; NADPH must bind to dNDUFA9 for CI assembly to proceed. Thus, forced expression of NDI1 in dIDH2-B thoraces restores the initial synthesis or stability of the Q-module.

**Figure 6:**
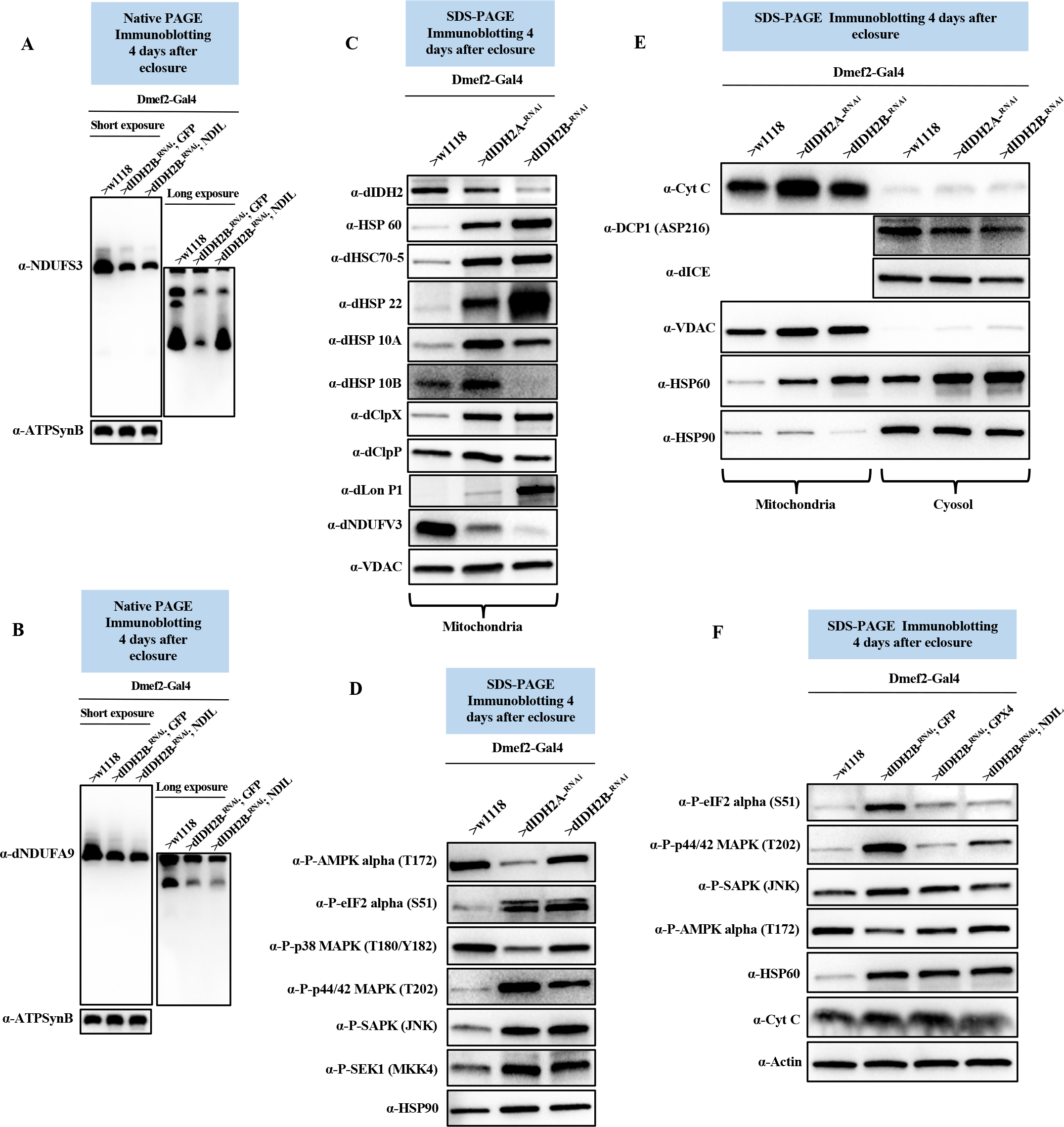
Induction of the alternative NADH-ubiquinone oxidoreductase (*Ndi1)* partially restores the biogenesis of the Q-module and suppresses the extent of activation of the JNK and MAPK pathways. (A and B) Mitochondrial preparations from thoraces isolated from Dmef2-Gal4/w1118 (wildtype), dIDH2-B flies overexpressing GFP (negative control), and dIDH2-B flies overexpressing ND1L four days after eclosure were analyzed by BN-PAGE, followed by immunoblotting with anti-NDUFS3 (A) and anti-dNDUFA9 (B). The blots were imaged following a short exposure to detect the holoenzyme and supercomplexes, after which the region corresponding to the holoenzyme and supercomplexes was cut off, and the rest of the blot re-imaged after a longer exposure to detect the assembly intermediates. Anti-ATP Synthase Beta (ATP5F1B) which detects the *Drosophila* ortholog, dATP-Syn*β* was used as a loading control. (C) Total cell lysates from flight/thoracic muscles isolated from Dmef2-Gal4/w1118 (wildtype), dIDH2-B and dIDH2-A flies 4 days after eclosure, were analyzed by SDS-PAGE and immunoblotting with the antibodies indicated; most of which detect mitochondrial chaperones and proteases. Anti-VDAC was used as a loading control. (D) Total cell lysates from flight/thoracic muscles isolated from Dmef2-Gal4/w1118 (wildtype), dIDH2-B and dIDH2-A flies 4 days after eclosure, were analyzed by SDS-PAGE and immunoblotting with the antibodies indicated; most of which detect stress-activated protein kinases. Anti-Hsp90 was used as a loading control. (E) Mitochondrial preparations (left three lanes) and cytosolic fractions (right three lanes) from flight/thoracic muscles isolated from Dmef2-Gal4/w1118 (wildtype), dIDH2-B and dIDH2-A flies 4 days after eclosure, were analyzed by SDS-PAGE and immunoblotting with the antibodies indicated. Anti-VDAC and anti-Hsp90 were used as loading controls for the mitochondrial and cytosolic fractions respectively. (F) Total cell lysates from flight/thoracic muscles isolated from Dmef2-Gal4/w1118 (wildtype), dIDH2-B flies overexpressing GFP (negative control), dIDH2-B flies overexpressing GPX4, and dIDH2-B flies overexpressing ND1L four days after eclosure were analyzed by SDS-PAGE and immunoblotting with the antibodies indicated. Anti-actin was used as a loading control.

To explore the effect of overexpressing NDI1 on the mitochondrial stress signaling network activated in dIDH2-A flight muscles, we first characterized the mitochondrial stress signaling cascade induced as a result of disrupting IDH2. Mitochondrial dysfunction activates a conserved stress signaling cascade termed the mitochondrial unfolded protein response (UPR^mt^) [28, 29] . The mitochondrial chaperones Hsp60, Mortalin, and Hsp10 as well as the Clp protease have all been associated with the UPR^mt^. The Clp protease consists of hexamers of a AAA+ ATPase (ClpX) and a tetradecameric peptidase (ClpP). In addition, Lon protease (Lon P1) is known to degrade oxidatively damaged proteins in the mitochondrial matrix [30]; and in *Drosophila* tissues, the mitochondrial chaperone Hsp22 is potently induced in response to mitochondrial distress [31]. Our proteomic studies described in Figure 5 suggested that an induction of genes that regulate the UPR^-mt^ and possibly other regulators of mitochondrial protein homeostasis may be a prominent component of the mitochondrial stress signaling network activated as a result of disrupting IDH2 (**Tables S1-S6**). Therefore, we generated antibodies to the Drosophila orthologs of Mortalin (Hsc70-5), Hsp10 (both paralogs), Hsp22, ClpP, ClpX and Lon P1 and used immunoblotting of proteins extracted from flight muscles to assess the relative expression of proteins implicated in mitochondrial protein homeostasis in dIDH2-A, dIDH2-B and wildtype thoraxes. Although dNDUFV3 expression was reduced – a reflection of the disintegration of CI – there was a potent upregulation of proteins that regulate the UPR^-mt^ (**Figure 6C**). These results indicate that disruption of dIDH2 enhances the UPR^-mt^, possibly to curb the damaging effects of enhanced ROS production on proteins.

When exposed to diverse stress stimuli, eukaryotic cells activate a common adaptive response referred to as the integrated stress response (ISR), to restore cellular homeostasis. A major event in this signaling cascade is the global downregulation of protein synthesis and the induction of a select set of cytoprotective genes, that act in unison to promote cellular recovery. Phosphorylation of the alpha subunit of eukaryotic initiation factor 2 (eIF2) is an established mechanism for limiting protein synthesis under various stress conditions [32]. Accordingly, we examined the extent of activation of various stress-responsive kinases and phospho-eIF2*α* (Ser51) (**Figure 6D**). There was an increase in the amount of phospho-eIF2*α*, phospho-p44/42 MAPK (Erk1/2), phospho-SAPK/JNK and phospho-SEK1/MKK4 in dIDH2-A and dIDH2-B thoraxes relative to wildtype samples. However, surprisingly, both phospho-AMPK and phospho-p38 were downregulated in dIDH2-A and dIDH2-B thoraxes (**Figure 6D**). Although the ISR is primarily an adaptive, pro-survival response, as the extent of the stress exposure increases, the signaling cascade can become maladaptive and ultimately induce cell death. Consequently, we examined whether cytochrome c was released from the mitochondria or whether caspases were cleaved, as an indication of the initial steps of apoptosis. In line with our proteomic results, cytochrome c levels were increased in the mitochondrion (**Figure 6E**). However, the upregulated cytochrome c was not released from the mitochondrion to cause apoptosis (**Figure 6E**); and we did not detect an increase in cleaved caspases (**Figure 6E**).

Finally, we examined the effect of upregulating NDI1 or the antioxidant enzyme (GPX4) on the stress signaling pathways activated in dIDH2-B thoraxes. Forced expression of NDI1 or GPX4 dampened the extent of activation of phospho-eIF2*a*, phospho-p44/42 MAPK (Erk1/2), and phospho-SAPK/JNK; and was associated with a relief-of inhibition of the phospho-AMPK pathway (**Figure 6F**). Nevertheless, the amount of Hsp60 or Cytochrome c was not appreciably altered (**Figure 6F**). Overall, we conclude from these results that the upregulation of cytochrome C in mitochondria from dIDH2-A and dIDH2-B thoraxes is not to promote apoptosis, but most likely promotes the transfer of electrons through a sub-optimal OXPHOS system.

### Disruption of dME3 produces a CI assembly intermediate profile that is similar to what is observed when dIDH2 is disrupted

Malic enzymes (ME) catalyze the oxidative decarboxylation of malate to pyruvate in a process that results in the reduction of NADP^+^ to NADPH, or NAD^+^ to NADH. In similarity to IDH, three ME isoforms exist. ME1 is localized to the cytosol while ME2 and ME3 are located in the mitochondrion. ME1 and ME3 use NADP^+^ as a cofactor. ME2 can use either NADP^+^ or NAD^+^ as a cofactor. CG5889 (Men-b) is the *Drosophila* ortholog of ME3. As is evident from Figure 1B, RNAi-mediated knockdown of dME3 causes a CI assembly defect. In view of the fact that both dME3 and dIDH2 reduce NADP^+^ to NADPH, and the OXPHOS assembly defect when dME3 is knocked down is similar to what was observed for IDH2-A, we wondered whether RNAi-mediated knockdown of dME3 in flight muscles using the Dmef2-Gal4 transgene (i.e. dME3-kd) produces a CI assembly intermediate profile that is similar to what is observed in IDH2-A thoraces.

Therefore, we examined the effect of disrupting dME3 on the biogenesis of the matrix domain of CI (**Figure 3A**). There was a reduction in the amount of dNDUFS3 in an initiating AI of the Q-module in mitochondria isolated from thoraxes of dME3-kd flies (**Figure 7A**). Similarly, the amount of dNDUFA9 that had incorporated into the Q-module was diminished in dME3-kd thoraces (**Figure 7B**). We also monitored the incorporation of dNDUFV1 and dNDUFV3 into CI AIs to ascertain the integrity of the N module. We observed that the stabilization or incorporation of both dNDUFV3 and dNDUFV1 into some AIs of the N-module were diminished in mitochondria isolated from thoraxes of dME3-kd flies (**Figures 7C and D**). This was coupled with a decrease in the amount of the Pp module subunits (dNDUFS5 and dNDUFA11) that had incorporated into subcomplexes from dME3-kd thoraces (**Figures 7E and 7F**). Importantly, incorporation of even the highly unstable mt-DNA encoded CI subunit, ND3, into a subcomplex in dME3-kd thoraces was not appreciably different from wildtype controls, indicating that a disruption of synthesis of mtDNA-encoded CI subunits is not the primary factor driving the CI assembly defect in dME3-kd flight muscles (**Figure 7G**). Finally, there was a slight stalling and accumulation of subunits in the PD module such as dNDUFB5 and dND5 (**Figure 7H**). Taken together, we conclude that disruption of dME3 impairs the biogenesis or stability of the Q and N-modules to trigger a disruption in CI assembly.

**Figure 7:**
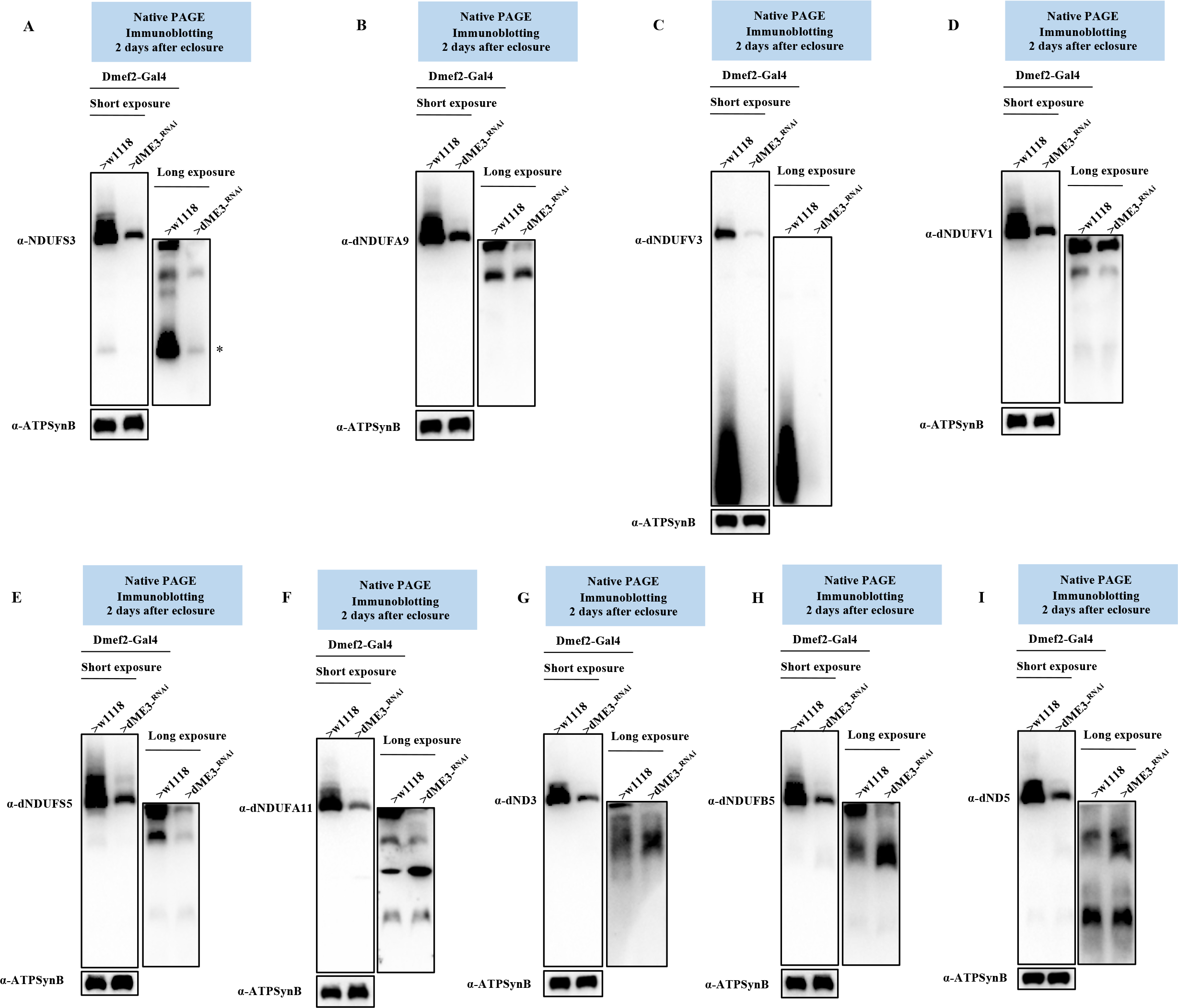
Disruption of dME3 produces a CI assembly intermediate profile that is similar to what is observed when dIDH2 is disrupted. (A-I) Mitochondrial preparations from thoraces isolated from Dmef2-Gal4/w1118 (wildtype) and Dmef2-Gal4/UAS-dME3^-RNAi^ flies 2 days after eclosure were analyzed by BN-PAGE, followed by immunoblotting with the antibodies listed. The blots were imaged following a short exposure to detect the holoenzyme and supercomplexes, after which the region corresponding to the holoenzyme and supercomplexes was cut off, and the rest of the blot re-imaged after a longer exposure to detect the assembly intermediates. The antibodies used were anti-NDUFS3 (A), anti- dNDUFA9 (B), anti-dNDUFV3 (C) anti-dNDUFV1 (D), anti-dNDUFS5 (E), anti-dNDUFA11 (F), anti-dND3 (G), dNDUFB5 (H), and anti-dND5 (I). Anti-ATP Synthase Beta (ATP5F1B) which detects the *Drosophila* ortholog, dATP-Syn*β* was used as a loading control.

## DISCUSSION

We have uncovered a signaling axis involving mitochondrial NADPH and ROS that links ferroptosis activation and OXPHOS assembly. We have found that when mitochondrial NADPH production is impaired as a result of dIDH2 disruption, ROS levels are elevated. This leads to a deterioration of the OXPHOS complexes in a defined order. It commences with a disruption of CI assembly, but progresses to the point where multiple OXPHOS complexes are impaired. NADPH is required for regenerating reduced glutathione in the glutathione cycle. Consequently, given that the ratio of oxidized glutathione to reduced glutathione increases when dIDH2 is knocked down, a plausible explanation for the increased ROS production is that insufficient amounts of reduced glutathione cause an increase in oxidative stress which ultimately destabilizes the OXPHOS complexes. Although the elevated ROS could in principle cause mutations in mtDNA to impair OXPHOS assembly, this is unlikely to be the primary driving force for the OXPHOS disintegration observed in dIDH2 flies; because we were able to detect AIs containing mtDNA-encoded CI subunits at a time when the matrix domain AIs were impaired (**see Figures 3 and 4)**. Thus, degeneration of the matrix domain precedes any purported damage to mtDNA.

The fact that CI is the first complex to disintegrate when dIDH2 is impaired suggests that this is unlikely to be due to non-specific ROS-induced protein oxidation, as non-specific protein oxidation would likely affect all complexes equally; precluding any specific and reproducible order of OXPHOS disintegration. This is especially true, given that knockdown of another mitochondrial NADPH-producing enzyme, dME3, also results in increased ROS production and a CI assembly defect. Accordingly, we propose that there are at least two signals that are triggered as a result of dIDH2 disruption to cause the specific order of OXPHOS degeneration observed. The first involves increased ROS levels; but the second is a decrease in the intramitochondrial pool of NADPH. A reduction in the intramitochondrial concentration of NADPH could in turn impact CI biogenesis in several ways. First, the interaction of NADPH with NDUFA9 is functionally significant, as it interacts with a conserved Arginine residue in NDUFS7, which contains the N2 Fe-S cluster [5]. Therefore, when the concentration of the intramitochondrial pool of NADPH is reduced, biogenesis of the Q-module will be stalled as well; because there will be a paucity of NADPH-containing NDUFA9 molecules. Furthermore, one of the reactions that occurs during the *de novo* synthesis of [2Fe-2S] clusters requires NADPH. Ferredoxin reductase (Arh1) is reduced by NADPH and then transfers its electrons to Yah1, where they are used for the synthesis of [2Fe-2S] on Isu1 (reviewed in [33]. As this is one of the initiating steps in the synthesis of both [2Fe-2S] and [4Fe-4S] clusters, a slight decrease in Arh1 reduction by NADPH will severely impair Fe-S cluster biogenesis. Indeed, the limited amount of intramitochondrial NADPH is likely the reason why although forced expression of ND1I restored biogenesis of an initiating Q-module subcomplex containing NDUFS3, it failed to fully ameliorate the Q module biogenesis defect.

When Fe-S cluster biogenesis is disturbed in the mitochondrial matrix, the amount of free intramitochondrial Iron increases. This can generate even more damaging species of ROS in the Fenton reaction and drive lipid peroxidation, ultimately resulting in ferroptosis. Ferroptosis is a non-apoptotic form of cell death that is triggered by iron-dependent lipid peroxidation [19]. Although Ferroptotic cell death has not been described in *Drosophila* flight muscles, several lines of evidence lead us to the conclusion that Ferroptosis is activated in dIDH2 thoraxes. First, although the lifespan of dIDH2-B flies is severely reduced, with most succumbing to lethality within 10 days of eclosure at 25C, caspase activity in the flight muscles of dIDH2-B flies was less than what is found in wildtype flies; indicating that apoptosis is not the primary cell death mechanism in dIDH2- B flies. Second, ROS, lipid peroxidation and the amount of labile (free) iron are all increased in the mitochondrion of dIDH2-B flies relative to wildtype controls. More importantly, raising the dIDH2 flies on two ferroptosis inhibitors – Ferrostatin-1 and Liproxstatin-1 – potently rescued their lethality. Finally, GPX4, is a Glutathione Peroxidase isoform that specifically reduces phospholipid hydroperoxides and oxidized lipoproteins to their native state, and several studies in mammalian systems have shown that disruption of GPX4 triggers lipid peroxidation and ferroptosis [19, 34]. Notably, we find that RNAi-mediated knockdown of the Drosophila ortholog of GPX4 also impairs OXPHOS assembly. Taken together, we conclude that induction of ferroptosis in flight muscles drives OXPHOS disintegration.

We note however that dIDH2 flies activate a robust compensatory stress signaling response that effectively buffers the flies against the pro-ferroptotic signals. As a case in point, while cytochrome c is robustly upregulated in the mitochondrion of thoraxes from dIDH2 flies, extramitochondrial cytochrome c pools were similar between wildtype and dIDH2 thoraxes, indicating that cytochrome c is not released into the cytosol to cause apoptosis. It appears the increased mitochondrial cytochrome c levels is an adaptive response that helps promote the transfer of electrons through a sub-optimally functioning electron transport chain. Similarly, our proteomic data revealed that several genes that regulate coenzyme Q biosynthesis were upregulated in dIDH2 samples. This result is also consistent with the induction of a robust adaptive response aimed at maintaining electron transfer through a sub-optimal ETC. Most notably, proteins that regulate the UPR^-mt^ and other aspects of mitochondrial protein homeostasis were strongly upregulated in dIDH2 thoraxes.

This may be one of the reasons why in spite of the induction of several pro-ferroptotic cues in dIDH2-A flies, neither their survival nor locomotory activity was impaired during the first two weeks after eclosure.

In summary, we have identified a pro-ferroptotic signaling network activated as a result of disrupting NADPH biosynthesis in the mitochondrion, that curtails OXPHOS assembly. This pro-ferroptotic signaling network is counteracted by a robust adaptive response that allows the flies to withstand the pro-ferroptotic cues. While a growing number of reports have shown that ferroptosis is implicated in multiple diseases such as cancer, neurodegeneration and ischemia/reperfusion injury in the heart, its role in skeletal muscle atrophy or sarcopenia is less clear. Nevertheless, given that skeletal muscles contain approximately 15% of the body’s total iron, and an age-dependent iron overload has been observed in skeletal muscles of rats, ferroptosis likely plays a role in muscle aging [35, 36]. We anticipate that future studies aimed at thoroughly defining the robust compensatory response induced in association with the pro-ferroptotic phenotype in dIDH2 flight muscles will be instrumental in unravelling additional mechanisms by which ferroptosis can be suppressed or induced.

## EXPERIMENTAL PROCEDURES

### *Drosophila* Stocks and Genetics

*Drosophila* stocks were reared at 25°C in vials containing agar, molasses, yeast and cornmeal medium supplemented with propionic acid and methylparaben in humidified incubators (Forma environmental chambers) on a 12-h:12-h dark: light cycle. Most of the transgenic stocks used were from the Bloomington Drosophila Stock Center (BDSC). GPX4 is what was previously described as GTPx-1 [37] The following transgenic stocks were used: *y w; Dmef2-Gal*, P{UAS-eGFP}34 (BDSC), P{UAS- Sod1.A}B.36 (BDSC), P{UAS-Sod2.M}UM83 (BDSC), P{UAS-Cat.A}2 (BDSC), UAS-GTPx and *UAS-NDI1.* The UAS-GTPx stock is referred to as UAS-GPX4 in this manuscript. RNAi stocks for disrupting Sod1/CG11793 were P{UAS-Sod1-IR}4 (BDSC), P{UAS-Sod1-IR}F103 (BDSC), P{TRIP.JF03321}attP2 (BDSC), P{TRIP.HMS00698}attP2 (BDSC), P{TRIP.HMS01291}attP2 (BDSC) and P{TRIP.GL01016}attP40 (BDSC); and were referred to as dSod1^-RNAi-1^ through dSod1^- RNAi-6^ respectively in Figure 1. RNAi stocks for disrupting Sod2/CG8905 were P{UAS- Sod2.dsRNA.K}15 (BDSC), P{TRIP.HMS00499}attP2 (BDSC), P{TRIP.HMS00783}attP2 (BDSC), and P{TRIP.GL01015}attP40 (BDSC); and were referred to as dSod2^-RNAi-1^ through dSod2^-RNAi-4^ respectively in Figure 1.

The following transgenic RNAi stocks were also used: Cat/CG6871 [P{TRIP.HMS00990}attP2 (BDSC)], dME-3/CG5889 [P{TRIP.HMC04802}attP40 (BDSC)], PRDX2/CG1633 [P{TRIP.HMS00501}attP2 (BDSC)], PRDX3/CG5826 [P{TRIP.HMJ22845}attP40 (BDSC)], PRDX4/CG1274 [P{TRIP.HMC04351}attP40 (BDSC)], PRDX5/CG7217 [P{TRIP.HMC05872}attP40 (BDSC)], PRDX6-1/CG3083 [P{TRIP.HMS05780}attP40 (BDSC)], PRDX6-2/CG12405 [P{TRIP.HMJ22929}attP40 (BDSC)], PRDX6-3/CG11765 [P{TRIP.GL00617}attP40 (BDSC)], GSTD1/CG10045 [P{TRIP.GL01039}attP2 (BDSC)], GSTD5/CG12242 [P{TRIP.HMS02534}attP40 (BDSC)], GSTS1/CG8938 [P{TRIP.HM05063}attP2 (BDSC)], Trxr-1/CG2151 [P{TRIP.HMJ21198}attP40 (BDSC)], and GPX4/CG12013 [P{TRIP.HMS00890}attP2 (BDSC)]. The transgenic RNAi stocks for disrupting dIDH2/CG7176 were P{GD6588}v42916 as dIDH2-A, P{KK107379}VIE-260B as dIDH2-B, and P{GD6588}v42915 as dIDH2-C, were from the Vienna *Drosophila* Resource Center (VDRC). P{TRIP.JF01989}attP2 (BDSC), P{TRIP.JF02173}attP2 (BDSC), P{TRIP.HMS00935}attP2 (BDSC), P{TRIP.HMJ23254}attP2 (BDSC) and P{TRIP.HMS05925}attP40 (BDSC) targeting Sod2, Cat, PRDX2, PRDX6-1 and GSTD1 respectively, were also tested for OXPHOS defects when crossed with Dmef2-Gal4, but were lethal.

### Locomotory Activity

Locomotory activity was assessed in two different ways.

Climbing Ability Assay: 20 adult male flies were placed in vials containing fly food. Subsequently, the vials were tapped gently to allow flies top settle at the bottom. The number of flies that climbed beyond the midpoint of the vial within 15 seconds were noted and recorded. This was repeated for 80 more flies (4 sets of 20). The number of flies that climbed beyond the midpoint of the vial were pooled together to obtain the number of flies that climb beyond the midpoint of the vial per 100 flies. The whole experiment was repeated 2 more times to obtain 3 biological replicates, representing a total of 300 flies Locomotory activity was assessed using the *Drosophila* activity monitor (TriKinetics). Specifically, 8 adult male flies were placed in the *Drosophila* activity monitor and spontaneous movements were recorded continuously for 288h on a 12-h:12-h dark: light cycle.

### Mitochondria Purification

Mitochondrial purification was performed essentially as described by Rera *et. al.* [12]. Fly thoraxes were quickly dissected and gently crushed with a dounce homogenizer (10 strokes) in 500μl of a pre-chilled mitochondrial isolation buffer (250mM sucrose, 0.15 mM MgCl_2_, 10mM Tris.HCl, pH 7.4) supplemented with Halt protease inhibitors (Pierce). Tissue homogenates were centrifuged twice, at 500g for 5 minutes at 4°C to remove the cuticle and other insoluble material. Subsequently, the supernatant was recovered and centrifuged at 5000g for 5 minutes at 4°C, to obtain the mitochondria-enriched pellet which was washed twice in the mitochondrial isolation buffer and stored at -80°C until further processing.

### Blue Native Polyacrylamide Gel Electrophoresis (BN-PAGE)

BN-PAGE was performed using NativePAGE gels from Life Technologies, and following the manufacturer’s protocol as previously described [11]. The digitonin: protein ratio used was 10g digitonin: 3g of protein.

### Silver Staining

Silver staining of native gels was performed with the SilverXpress staining kit from Life Technologies, following the manufacturer’s instructions.

### In-gel Complex I, II, IV and V Activity

In-gel Complex I activity was assessed by incubating the native gels in 0.1 mg/ml NADH, 2.5 mg/ml Nitrotetrazolium Blue Chloride (NTB), 5 mM Tris-HCl (pH 7.4) at room temperature.

In-gel Complex II activity was assessed by incubating the native gels in 20 mM sodium succinate, 0.2 mM phenazine methasulfate, 2.5 mg/ml NTB, 5 mM Tris-HCl (pH 7.4) at room temperature.

In-gel Complex IV activity was assessed by incubating the native gels in 50 mM sodium phosphate (pH 7.2), 0.05 % 3,3’- diaminobenzidine tetrahydrochloride (DAB), 50 μM horse heart cytochrome C at room temperature.

In-gel Complex V activity was assessed by pre-incubating the gel in 35 mM Tris-base, 0.27M glycine pH 8.4 for 3 hours and then subsequently in 35 mM Tris-base, 0.27M glycine (pH 8.4), 14 mM MgSO4, 0.2% w/v Pb(NO3)2 and 8 mM ATP at room temperature.

### Amplex Red Assay for Measuring Hydrogen Peroxide Production

The amount of hydrogen peroxide produced was monitored using the Amplex Red Hydrogen Peroxide/Peroxidase Assay Kit (Catalog no. A22188, Molecular Probes). In brief, fly thoraxes were homogenized in pre-chilled mitochondrial isolation buffer supplemented with halt protease inhibitors (Pierce), and centrifuged twice at 500g for 5 minutes at 4°C to remove insoluble material. Subsequently, a serial dilution of the supernatant was added to a reaction buffer containing 100uM Amplex Red Reagent and 0.2U/ml Horseradish Peroxidase solution. Fluorescence was measured at an excitation wavelength of 540nm and detected at 590nm every 30 sec for 30 min at 25°C using a SpectraMax paradigm multi-mode microplate reader (Molecular Devices). The background fluorescence determined for a no H2O2 reaction was deducted from each value. Amplex Red activity was normalized to protein concentrations as determined with a microBCA kit (Thermo Fisher Scientific, Waltham, MA).

### Lipid Peroxidation Assay

The extent of lipid peroxidation in fly thoraces was assessed using the lipid peroxidation assay kit (Sigma catalog number MAK085). 30 fly thoraxes were homogenized in 0.3 mL of MDA lysis buffer and 3 μL of 100X BHT; both solvents were supplied with the kit. The samples were subsequently centrifuged at 13,000g for 10 minutes to remove the cuticle and other insoluble material. After protein concentrations were normalized between samples, 200 μL of each sample was aliquoted into separate eppendorf tubes. Subsequently, 200 μL of Perchloric acid was added to each tube, vortexed and centrifuged at 13,000 g for 10 min. 200 μL of the recovered supernatant was transferred into a new eppendorf tube, and 600 μL of the TBA solution (supplied with the kit) was added to form the MDA-TBA adduct. The samples were subsequently incubated at 95 ^0^C for 60 minutes, and then placed in an ice bath for 10 minutes. To enhance sensitivity, 300 μL of 1-butanol was added to extract the MDA-TBA adduct from the 800 μL reaction mixture, and then centrifuged at 16,000g for 3 minutes at room temperature to separate the layers. The fraction containing 1- butanol (i.e. the top layer) was transferred to another tube. Following evaporation of the 1-butanol, the residue containing the MDA-TBA adduct was dissolved in 200 μL of water, and then transferred to a 96 well plate. Absorbance of the MDA-TBA adduct was measured at 532 nm; and the amount of MDA present in duplicate samples assessed using a MDA standard graph.

### NADP/NADPH Quantification

The NADP/NADPH ratio was measured in fly thoraces with the NADP/NADPH Quantification kit (Sigma # MAK038). Briefly, 10 fly thoraxes were homogenized in 500 μL of NADP/NADPH Extraction Buffer (supplied with the kit) and 5 μL of 100X Halt protease inhibitor (Pierce); and the samples centrifuged at 10,000g for 10 minutes to remove insoluble material. Any residual particulate matter was removed by transferring the extracted NADP/NADPH supernatant into a 10 kDa cut-off spin filter; and centrifuging at 10,000g for 20 minutes at 4^0^ C. The flow-through (extracted samples) was saved for further analyses.

To detect NADPH, NADP was decomposed by aliquoting 200 μL of the extracted samples into Eppendorf tubes; and heating to 60^0^ C for 30 minutes in a heating block. Samples were cooled on ice.

50 μL of both the decomposed NADP (measures NADPH only) and regular samples (measures both NADPH and NADP) were aliquoted into 96-well plates in duplicate; and 100 μL of the Reaction Mix supplied with kit was added to each of the standard and sample wells. The samples were mixed thoroughly, and incubated for 5 minutes at room temperature, to convert NADP to NADPH. 10 μL of NADPH developer (supplied with the kit) was added to each sample and incubated at room temperature for 60 minutes, after which the absorbance of both samples was measured at 450 nm. Subsequently, the reaction was stopped by adding 10 μL of Stop Solution (supplied with the kit) to each well and mixing thoroughly. The background value for the assays was the value obtained for the blank NADPH standard. After correcting for the background value the amount of NADPH present in the sample was assessed using the NADPH standard curve; and the NADP/NADPH ratio by using the formula.

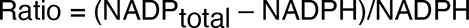

### Mitochondrial free Fe^2+^ measurement

The mitochondrial free Fe^2+^ was measured using the QuantiChrom Iron Assay Kit. Briefly, mitochondria from 40 thoraxes were suspended in 300 μL of medium containing 5 mM PIPES buffer, pH 6.5 and incubated at 37° C for 30 minutes to produce a hypo-osmotic swelling. The ruptured mitochondrial sample was subsequently centrifuged at 10,000g for 5 minutes; after which the supernatant was recovered. 200 μL of the working reagent (supplied with the kit) was added the absorbance was measured at 590 nm. The background value for the assay was the value obtained for the blank used together with the varying concentrations of Fe to plot the standard curve. Samples were analyzed in triplicates. After correcting for background values, the amount of Fe^2+^ present in the sample was estimated using the Fe standard curve.

### Caspase activity assay

Caspase 3 activity was measured by using a Caspase 3 Fluorimetric Assay Kit **(Sigma # CASP3F)**. In brief, fly thoraxes (10 thoraxes) were homogenized in 1X PBS supplemented with halt protease inhibitors (Pierce), and centrifuged twice at 500g for 5 minutes at 4°C to remove insoluble material. 200 μL of the Reaction mixture (supplied with the kit) containing 250 uM Caspase 3 substrate (Ac-DEVD-AMC) was added to each of the standards and samples, thoroughly mixed, and incubated for 60 minutes at room temperature, protected from light. Fluorescence was measured at an excitation wavelength of 360 nm and detected at 460 nm every 30 sec for 60 min at 25^0^ C using a SpectraMax microplate reader. After correcting for background values, caspase 3 activity was assessed using the AMC standard curve.

### Ferroptosis rescue

Flies of the appropriate genotype were allowed to lay eggs for 48 h at 25° C. After the ensuing larvae reached the third instar, 200 uL each, of 1 mg/mL Ferrostatin-1 and 1 mg/mL Liproxstatin- 1, were added to the fly food; and the larvae were allowed to continue developing in the midst of the drug until they eclosed. The adult flies were reared at a density of 25 flies per vial; and maintained at 25° C. Flies were transferred to vials of fresh fly food every other day and scored for viability every day.

### Generation of Peptide Polyclonal Antibodies

Rabbit polyclonal antibodies recognizing various segments of specific target proteins in *Drosophila* were generated by Biomatik using the synthetic peptides listed below:

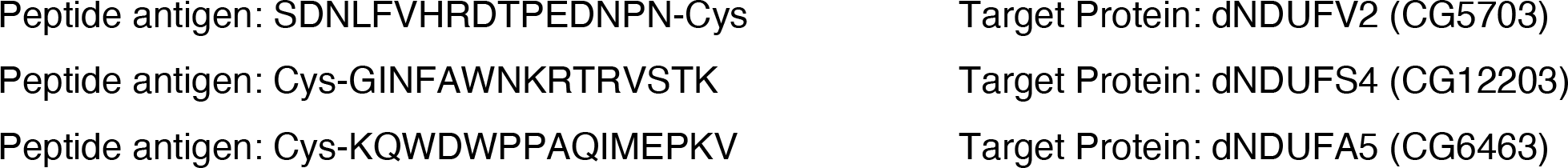
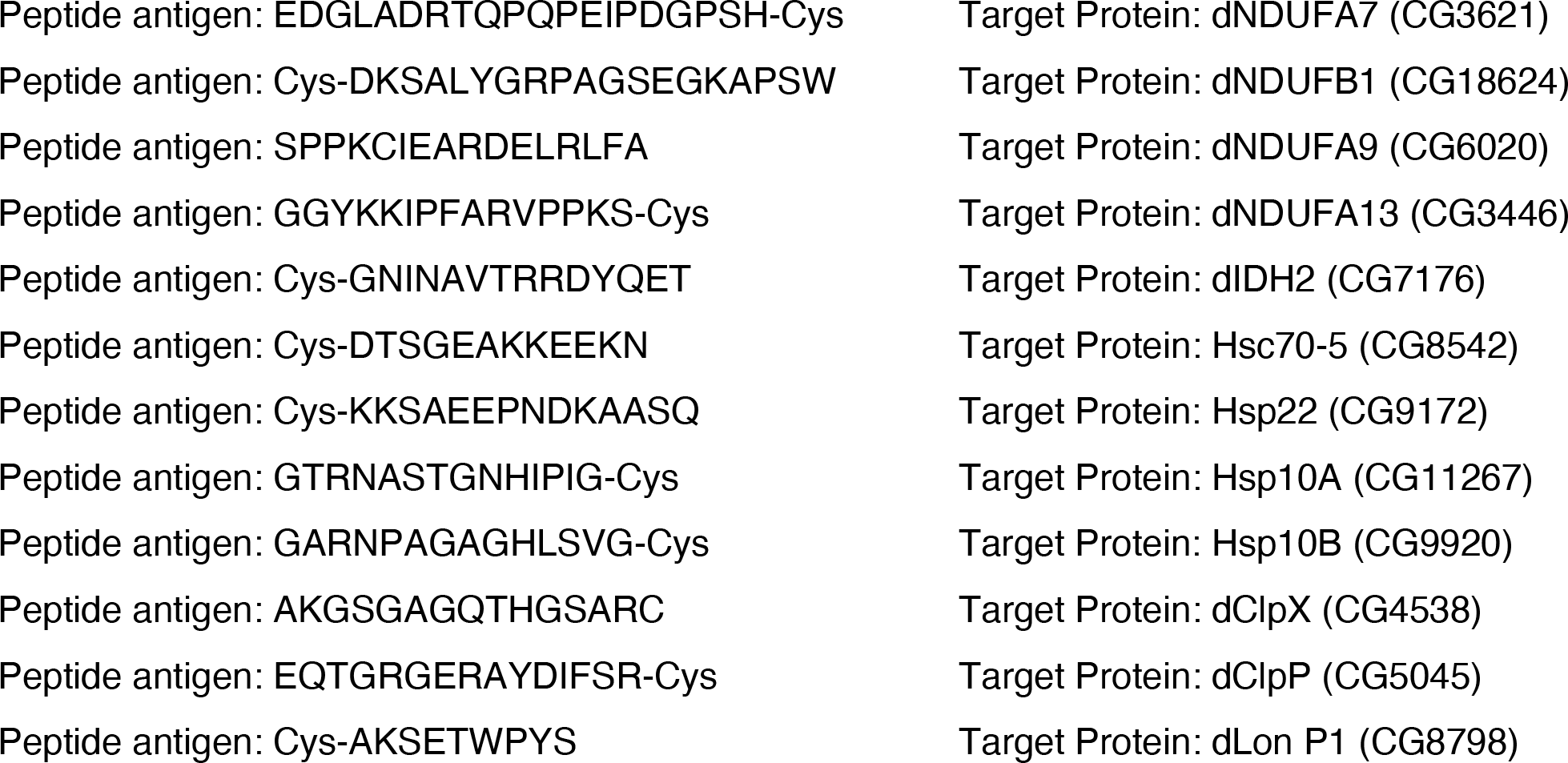

### Immunoblotting

Immunoblotting was performed as previously described [11]. In addition to the new rabbit polyclonal antibodies we generated, the following primary antibodies were also used: anti- dNDUFS2 [11], anti-NDUFS3 (abcam, ab14711), anti-dNDUFS5 [11], anti-dNDUFS7 *(Murari et. al. 2021),* anti-dNDUFS8 *(Murari et. al. 2021),* anti-dNDUFA8 *(Murari et. al. 2021),* anti-dNDUFA11 *(Murari et. al. 2021),* anti-dNDUFA12 *(Murari et. al. 2021),* anti-dNDUFV1[11], anti-dNDUFV3 *(Murari et. al. 2021),* anti-dNDUFB5 [11], anti-dNDUFB6 [11], anti-dNDUFB8 [11], anti-dND1 [11], anti-dND2 [11], anti-dND3 [11], anti-dND4L [11], anti-dND5 [11], anti-dND6 [11], anti-VDAC (abcam, ab14734), and anti-ATPsynß (Life technologies, A21351). Secondary antibodies used were goat anti-rabbit Horseradish Peroxidase (PI31460 from Pierce) and goat anti-mouse Horseradish Peroxidase (PI31430 from Pierce).

A commercially available marker [i.e. the NativeMark Protein standard (Life Technologies)], which is a soluble protein marker, was initially used to estimate the molecular weight of the protein complexes. However, there are major discrepancies between the migration behavior of membrane and soluble protein markers [38]. Accordingly, estimating the sizes of membrane proteins such as OXPHOS complexes or assembly intermediates on blue native gels using standard soluble protein markers can be problematic. Due to these discrepancies, where possible, the identity of OXPHOS complexes and assembly intermediates in immunoblots were assessed based on their position in the gel and known constituent protein subunits.

### In-gel Digestion and Mass Spectrometry

OXPHOS complexes from mitochondrial preparations from Dmef2-Gal4/w1118 (wildtype) and Dmef2-Gal4/UAS-dIDH2A^-RNAi^ flies were separated on blue native gels. Each sample was run in triplicate. Following resolution of the OXPHOS complexes, the gel was incubated in a fixative consisting of 50% methanol, 10% acetic acid and 100mM ammonium acetate for 30 minutes. Subsequently, the gel was washed twice with ultrapure water, and six gel slices corresponding to the holoenzyme, and 5 slices between the region where CIV and CII migrate were excised for each genotype. The excised gel slices were further diced into smaller pieces, placed in eppendorf tubes, and de-stained in a gel de-staining buffer (8% Acetic acid). In-gel trypsin digestion was performed essentially as described previously [39]. In brief, 100µl of 25mM DTT and 100µl of 50mM iodoacetamide were used for protein reduction and alkylation respectively, followed by digestion with 0.5 µg of trypsin at 37 °C for 16 h. The tryptic peptides were extracted and desalted using a C18 cartridge, followed by LC-MS/MS analysis on an Orbitrap Fusion Lumos Tribrid mass spectrometer (MS) (Thermo Scientific). The peptides were separated on a C18 nano column (75 μm × 50 cm, two μm, 100 Å) with a 2-h linear-gradient consisting of solvent A (2% acetonitrile in 0.1% formic acid (FA)) and solvent B (85% acetonitrile in 0.1% FA). The eluted peptides were directly introduced to the MS *via* a nanospray Flex ion source. The MS spectra were acquired in the positive mode with a spray voltage of 2 kV. The temperature of the ion transfer tube was 275°C. MS scan range was between m/z 375 and 1,500 with a 120,000 (FWHM) resolution in the Orbitrap MS. The peptides with charge states between 2 and 7 were selected for MS/MS analysis. Higher-energy Collisional Dissociation (HCD) was used for peptide fragmentation with the collision energy of 30%.

### Protein Identification and Quantification

The MS/MS spectra were searched against the Uniprot *Drosophila* database (21,107 entries, downloaded on 11/12/2020) using the Sequest search engine through the Proteome Discoverer (version 2.4) platform. The mass tolerance was 10 ppm for MS and 0.6 Da for MS/MS. Methionine oxidation and N-terminus acetylation were set as variable modifications, and cysteine carbamidomethylation was set as a fixed modification. The false discovery rate accepted for the identification of both proteins and peptides was less than 1%. Relative protein quantitation was calculated based on the spectral counting (SC) method [24, 40]. To circumvent the problem with large SC ratios from small spectra counts in the SC ratio denominators, we arbitrarily added 2 SC for each protein before calculating the protein quantification ratios.

### Statistics

Except where noted, p values are based on the student’s t-test for unpaired two-tailed samples. The fold change shown refers to the mean ± s.e.m (standard error of the mean); and * = p<0.05, ** = p<0.01 and *** = p<0.001.

## Acknowledgements

We thank past and present members of the Owusu-Ansah lab for general discussions; Henry Colecraft, Jeanine D’Armiento, Alexander Galkin, Wes Grueber, Laura Johnston, Andrew Marks, Martin Picard, Liza Pon, Michael Schlame, Eric Schon and Mimi Shirasu-Hiza for fly stocks, reagents, and critical discussions. We acknowledge the Bloomington *Drosophila* Stock Center and the Vienna *Drosophila* Resource Center for various fly strains. The mass spectrometry data were obtained from an Orbitrap mass spectrometer funded in part by NIH grants NS046593 and 1S10OD025047-01, for the support of proteomics research at Rutgers Newark campus. Work in the Owusu-Ansah lab is supported by a Provost Junior Faculty grant to support Columbia’s diversity efforts, and NIH grants DK112074 (R21), AR077312 (R21), and GM124717 (R35).

## Author contributions

E. Owusu-Ansah conceived the project, designed all experiments, and supervised the work. A. Murari, S. K. Rhooms, K.B.F. Hossain, N.S. Goparaju, C. Osei and E. Owusu-Ansah performed all experiments except the acquisition of the mass spectrometry data, which were performed by T. Liu and H. Li. A. Murari and E. Owusu-Ansah analyzed and discussed results. T. Liu and H. Li wrote the proteomics section of the methodology and limitations of the study. E. Owusu-Ansah wrote all other sections of the manuscript, and got feedback from A. Murari.

